# Sildenafil-driven cone PDE6 inhibition alters receptive-field properties of retinal ganglion cells ex vivo

**DOI:** 10.1101/2025.10.07.680926

**Authors:** Bartłomiej Bałamut, Piotr Węgrzyn, Anna Posłuszny, Andrzej T. Foik

**Affiliations:** International Centre for Translational Eye Research (ICTER), Institute of Physical Chemistry PAS, Marcin Kasprzak St., 01-244 Warsaw, Poland; Institute of Physical Chemistry, PAS, Marcin Kasprzak St., 01-244 Warsaw, Poland; University of Warsaw, Faculty of Physics, Pasteur 5, 02-093 Warsaw, Poland

**Keywords:** phototransduction, *ex vivo* recordings, retina, retinal degeneration, visually evoked responses

## Abstract

Suppressing the phototransduction cascade has a profound impact on visual information processing in the retina. Here, we examined how acute silencing of photoreceptors alters stimulus preference of Retinal Ganglion Cells (RGCs) without affecting retinal anatomy. Using ex vivo recordings from C57BL/6 mouse retinas, we applied the phosphodiesterase 6 (PDE6) inhibitor Sildenafil and compared responses to flash and drifting-grating stimuli before and after treatment.

Flash responses revealed six physiological RGC populations under control conditions. After Sildenafil, Off-responses were completely abolished, and most cells adopted On-like profiles with markedly prolonged response latencies. At the population level, orientation selectivity was preserved; however, RGCs lost the ability to detect spatial frequencies above 0.05 cycle/° and temporal frequencies above 2 cycle/s. A subset of cells increased their firing rates and transformed from orientation-selective into direction-selective cells, indicating reorganization of tuning properties.

To assess photoreceptor specificity of Sildenafil, we tested Gnat1 (rod-deficient, cone-only) and Gnat2 (cone-deficient, rod-only) mouse models. Sildenafil completely removed visually evoked responses in Gnat1 retinas, but not in Gnat2, demonstrating preferential inhibition of cone PDE6.

These results show that cone-selective suppression of phototransduction is sufficient to abolish Off-pathway signaling, reduce high-frequency visual sensitivity, and alter RGC receptive-field tuning. Beyond providing a mechanistic explanation for the visual side effects of PDE5/6 inhibitors, this establishes a pharmacological model to mimic early cone dysfunction and study inner retinal adaptation.

**Significance Statement:** Photoreceptor degeneration alters visual processing long before anatomical changes are evident, but the early functional consequences for retinal circuits remain poorly defined. Here, we show that pharmacological suppression of phototransduction with the PDE6 inhibitor Sildenafil selectively abolishes cone-driven responses and profoundly alters retinal ganglion cell tuning. Off-pathway signaling is lost, spatiotemporal frequency sensitivity is reduced, and some cells acquire direction selectivity. These findings provide functional evidence that cone pathways are preferentially targeted by Sildenafil, offering a mechanistic explanation for its transient visual side effects in humans. More broadly, this approach establishes a pharmacological model for studying inner retinal adaptation to early cone dysfunction in degenerative disease.

## Introduction

Mice’s retina efficiently processes visual information and, therefore, can encode complex features of visual scenes, such as orientation selectivity, contrast discrimination, and spatial and temporal frequencies (Baden et al., 2016; Ding et al., 2016; Reinhard and Münch, 2021; Kerschensteiner, 2022; Kim et al., 2022; Strauss et al., 2022; Barta and Thoreson, 2024; Hsiang et al., 2024). Therefore, any change in photoreceptor activity has a ripple effect on the entire retina and the entire visual system (Niell, 2013; Chen et al., 2016; Rhim et al., 2017, 2021; Rasmussen et al., 2020). Unfortunately, the physiological changes are usually visible only after the retina is severely degenerated. Negative plasticity progresses simultaneously with the initial physiological changes of the retinal ganglion cells (RGCs), which occur when photoreceptors start dying or becoming nonfunctional. Investigating the early changes in activity of the RGCs at the time when photoreceptor degeneration begins may help to identify noninvasive markers of early retinal degeneration and to develop indicators for properly restored vision (Arenkiel et al., 2007; Thyagarajan et al., 2010).

We aimed to characterize the activity of RGCs after partial suppression of the phototransduction cascade using Sildenafil, a known inhibitor of phosphodiesterase 6 (PDE6) (Tomczewski et al., 2025). Using a high dose of Sildenafil enabled us to observe changes happening immediately after the inhibitor was applied to the *ex vivo* mouse retina. The most visible effect was related to the decrease of the evoked-potential amplitudes and single-cell light responses. We investigated how suppression of photoreceptors changes the responses of RGCs to drifting grating, similar to responses recorded in various rodent models previously (Foik et al., 2015, 2018, 2020; Suh et al., 2020; Frankowski et al., 2021; Choi et al., 2022; Leinonen et al., 2022; Lewandowski et al., 2022; Xu et al., 2022; Engfer et al., 2023). In the healthy state of retinas from control mice, we found broad spatial and temporal frequency tuning and orientation selectivity. In the retinas from drug-treated mice, those parameters changed towards slower and wider grating preferences; moreover, a subset of cells increased their firing rate and changed physiological features towards different physiological groups.

## Methods

### Animals

All experiments with mice (C57Bl/6J) followed the European Union directive on laboratory animal use (2010/63/EU) to minimize pain and suffering, and adhered to Polish laws about the use of laboratory animals.

### Retina preparation

Mice were overdosed with isoflurane. After confirmation of no response to pinching of their feet, mice were decapitated with decapitation scissors. Eyes were removed with Vannas scissors and submerged in Ames-HEPES solution (adjusted to pH 7.4) (Fujii et al., 2016; Rathbun et al., 2024). The cornea of each eye was punctured with an 18-G needle and subsequently removed with microsurgical scissors. Then, the lens was removed, and the eyecup was transferred to a fresh dish. Both retinas were removed from the eyecups and placed in fresh Ames-HEPES solution. The vitreous was mechanically removed from each eye using two forceps. The retina was cut into a “clover shape” and placed on nitrocellulose filter paper with a puncture in the middle. Such a construct was placed on a flat petri-dish cover, and the retina was further straightened before being placed in an MEA chamber with the retinal ganglion cells facing the contacts. The retina was clamped to the bottom of the dish with a 1-gram horseshoe clamp (Fujii et al., 2016).

### Multi-electrode array (MEA) recordings

For MEA-recordings, standard chips (60MEA100/10iR-Ti) were used, manufactured by Multi-Channel Systems (Germany). Recordings were carried out in the MEA2100-Mini system with a built-in heating plate. The recording chamber was constantly perfused with carbonated Ames-bicarbonate solution (4 ml/min). Each retina was incubated for 30-40 minutes before recordings were performed (Figure 1A). Data were collected using dedicated software provided by the producer (Multi Channel Systems), and later analyzed with custom-written MATLAB (MathWorks, USA) scripts. Raw data were sampled at 40 kHz and then split into low-frequency local-field-potential (LFP) data from 0.1 Hz to 300 Hz, and high-frequency data from 0.3 to 7 kHz, containing action potentials (spikes). The low-pass signal was aligned to stimulus time stamps, down- sampled to 1kHz, and averaged across trials to calculate visually evoked potentials (VEPs).

**Figure 1.**
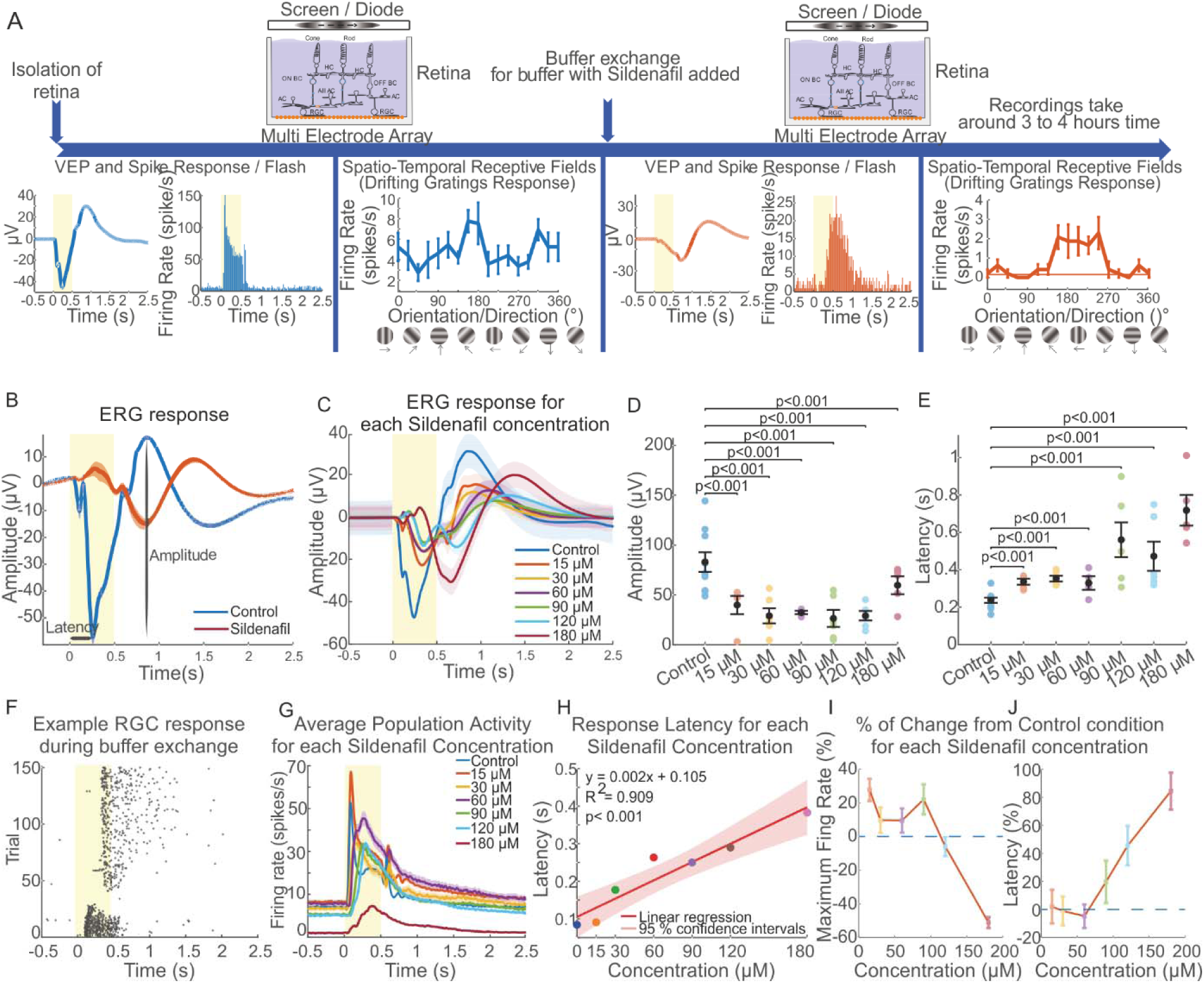
Suppression of phototransduction changes visually evoked activity. A, Experimental workflow of the *ex vivo* retinal recordings in the control condition and after the addition of Sildenafil to suppress the phototransduction cascade. B, Population-averaged VEP to a 0.5 s flash of light in the control medium (blue), and with added Sildenafil (red). C, Population VEPs for the Control and each Sildenafil concentration tested in the experiment. Concentrations are indicated in the panel legend and encoded by color. D, Population summary showing a comparison of the amplitude responses before (blue), and after Sildenafil addition at different concentrations (encoded by color). E, Comparison of the population-response latency to the flash stimulus before (blue), and after the Sildenafil addition, as in D. F Changes in the delay of single-cell responses to a 0.5-s flash of light in subsequent trials recorded during the exchange of buffer (in the first minutes from the introduction of the buffer containing Sildenafil). G, the population average response histograms for all used concentrations encoded by different colors. H, the linear trend of the response latency related to the Sildenafil concentration. I, the relative change of the population maximum firing rate vs. the inhibitor concentration. J, The relative change in response latency vs. the drug concentration.

Visual stimuli were generated in MATLAB using the Psychophysics Toolbox and displayed on a gamma-corrected LCD monitor (Waveshare 12030, China), focused on the photoreceptor layer to minimize distortion and ensure precise mapping of spatial frequencies. The optical relay was designed as a 4F system built with achromatic doublets (to correct for chromatic and spherical aberrations). Using lens L1 (focal length 400 mm, Thorlabs AC254-100-A) and lens L2 (focal length 30 mm, Thorlabs AC254- 050-A), we achieved a 0.075 demagnification, covering the full area of the multi-electrode array (MEA; 2 × 2 mm) with uniform illumination and minimal distortion. Stimulus-onset times were corrected for monitor delay using an in-house designed photodiode system (Foik et al., 2018, 2020). For visually evoked response recordings and cell type classification, retinas were tested with 50 repetitions of a 500-ms mesopic flash of light (1.5 photopic cd/m²) (Zele and Cao, 2015). Receptive fields (RFs) for visually responsive cells were then defined using square-wave drifting gratings, after which optimal orientation/direction and spatial and temporal frequencies were determined using sine-wave gratings. Spatial frequencies (SFs) tested were from 0.001 to 0.5 cycles/°. Temporal frequencies (TFs) tested were from 0.1 to 9 cycles/s. With the optimal temporal and spatial frequency, and at high contrast, the orientation tuning of the cell was tested again using 8 orientations at 2 directions each, stepped by 22.5° increments. After all these tests, the regular buffer was exchanged for a buffer containing Sildenafil (Maxon Forte, Adamed, Poland) at a concentration of 0.087 mg/ml (180 µM) (Martins et al., 2015). In the first 10 minutes of exposure to Sildenafil citrate, the retina was stimulated with 500 ms flash every 10 seconds. Afterward, the spatiotemporal RF test procedure was repeated to assess changes in response parameters of RGCs induced by Sildenafil.

### Data analysis

All recordings were submitted to Offline spike sorting with the wavelet-sorting algorithm using the WaveClus toolbox (Chaure et al., 2018).

Cells were classified into six groups based on the shape and component composition of the response to a 500 ms flash of light: On Sustained, Off Sustained, On Transient, Off Transient, On-Off, or “Other”. The “Other” category was used for cells that could not be classified into any of the other designated groups. When cells showed visually driven activity, we used further drifting grating stimuli to assess the properties of the spatiotemporal receptive fields. Each cell was evaluated for orientation selectivity, optimal spatial frequency, and optimal temporal frequency. A cell was categorized as grating responding if it was possible to determine one of the above-mentioned optimal parameters describing the receptive field.

Tuning curves were calculated based on the average firing rate. Optimal visual parameters were chosen as the maximum response value. Orientation tuning was measured in degrees at the Half-Width at Half-Height (HWHH; 1.18 x σ) based on fits to a double Gaussian function (Foik et al., 2018):

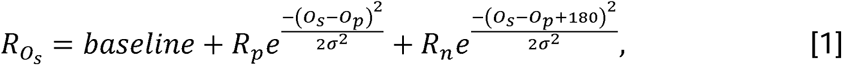

where O_s_ is the stimulus orientation, R_Os_ is the response to different orientations, O_p_ is the preferred orientation, R_p_ and R_n_ are the responses at the preferred and non-preferred directions, σ is the tuning width, and ‘baseline’ is the offset of the Gaussian distribution. Gaussian fits were estimated without subtracting spontaneous activity.

The optimal spatial and temporal frequency was extracted from the data fitted to the Gaussian function using equation [2] (Foik et al., 2018, 2020):

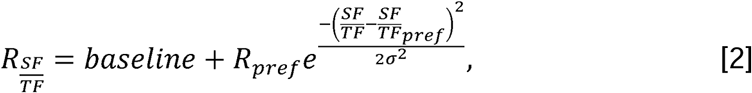

where R_SF/TF_ is the estimated response, and R_pref_ indicates the response at a preferred spatial or temporal frequency. SF/TF indicates spatial or temporal frequency, σ is the standard deviation of the Gaussian, and baseline is the Gaussian offset.

### Statistical Analysis

Data throughout the manuscript are presented as mean ± standard error of the mean (SEM). The level of statistical significance was set at P < 0.05 for nonparametric two-tailed Mann-Whitney or Wilcoxon U-tests. Offline data analysis and statistics were performed in MATLAB.

## Results

### Sildenafil concentration-dependent activity of RGCs

The C57B6 retinas were persisting in activity preservation after adding the phototransduction cascade inhibitor – Sildenafil. Thus, we decided to prepare an activity tuning curve to investigate in detail how Sildenafil changes Retinal Ganglion Cells (RGCs) activity. Interestingly, we have noticed that the process is not entirely linear, but rather has a dramatic influence on electrical activity after crossing 90 µM Sildenafil concentration. Overall, the net firing rate activity of single cells increased from 20 to 40 % over the baseline; however, when the concentration was higher than 90 µM, the activity became negative, eventually reaching -55% change from the control condition (Figure 1). The only close to linear effect we observed was for the response latency, which almost consistently increased with the sildenafil concentration. Another interesting observation was that for the 180 µM, the visually evoked potentials (VEP) amplitude did not differ significantly from the control condition, even if all the components were delayed in time by 300 ms. Having these findings, we decided to use the 180 µM Sildenafil concentration to test its influence on the drifting grating processing in the retina after phototransduction is silenced.

### High concentration of Sildenafil affects motion processing in wild-type retinas

In all of the tested retinas (n = 5), we found robust and strong visually evoked potentials (VEPs) and single-cell activity (Figure 1A), with the number of classified cells presented in Table 1. Initially, we tested retinal responses to screen flashes to assess the strength of the responses. Then we tested all single cells for the drifting-gratings optimal parameters. After reference recordings, we replaced the medium with fresh medium containing the phosphodiesterase inhibitor - Sildenafil, and all of the stimuli were tested again. After the Sildenafil addition, we found that the shape of the VEPs reversed (Figure 1B), and the amplitude of the responses was significantly decreased. We observed changes in the shape and magnitude of the response with the lowest concentration used, and progressed with increasing concentration. However, it never entirely turned off the response. Therefore, this convinced us to test drifting grating responses with the highest concentration of Sildenafil. In the control condition, the average population VEP amplitude was 80.28 ± 4.27 µV; after suppressing the phototransduction cascade with the PDE6 inhibitor, the amplitude dropped to 35.68 ± 2.83 µV (P < 0.001; Figure 1D). Even though the drop in the response amplitude and reversal of the potential were very prominent, the most striking result was the change in response latency. The latency of the most negative components of the response in the control condition was 238.0 ± 0.6 ms, whereas after Sildenafil addition, it increased to 815.00 ± 0.02 ms (P < 0.001; Figure 1E). This extension of the response latency was strictly related to the Sildenafil concentration, as we present in the plots. We did not observe a full restoration of evoked activity after the washout of Sildenafil. Similar changes are also reflected in the activity of single cells. In Figure 1F, we present an example of the activity of a single RGC while Sildenafil was in the recording chamber. In the first phase, the activity dropped significantly, and the cell changed its response pattern with the appearance of persistent activity after stimulus cessation. The subset of cells became completely silent for several trials, as presented in the example in Figure 1F, and most of the cells smoothly transitioned to very long response latencies or were attenuated. Again, as in the VEPs analysis, we did not observe the entire silencing of the flash-evoked response in single RGCs (Figure 1G). However, we discovered that both the response intensity (firing rate) and response latency (Figure 1H) in a linear trend. Interestingly, for initial concentrations, the population maximum firing rate was higher than in the control condition and dropped only above 120 µM (Figure 1I). The response latency was delayed starting from 90 µM concentration of the inhibitor (Figure 1J).

**Table 1.** Summary of the number of cells recorded before and after Sildenafil addition.

|  |  | Flash<br>Response | Grating<br>Response |
| --- | --- | --- | --- |
| Control | On<br>Sustained | 100 | 79 |
|  | On<br>Transient | 87 | 49 |
|  | Off<br>Sustained | 8 | 7 |
|  | Off<br>Transient | 22 | 10 |
|  | On-Off | 141 | 100 |
|  | Other | 38 | 17 |
|  | Total: | 396 | 262 |
| Sildenafil | On<br>Sustained | 92 | 71 |
|  | On<br>Transient | 51 | 13 |
|  | Off<br>Sustained | 7 | 6 |
|  | Off<br>Transient | 13 | 1 |
|  | On-Off | 120 | 79 |
|  | Other | 34 | 17 |
|  | Total: | 317 | 187 |

To analyze RGC responses in detail and compare between 0 and 180 µM concentration of PDE6 inhibitor, we decided to divide the populations according to the basic properties of their flash responses. Accordingly, we distinguished six cell populations: Off-Sustained, Off-Transient, On-Off, On-Sustained, On-Transient, and Other. For the “Other” group, we included all cells that could not be assigned to any of the designated types (Goetz et al., 2022).

In Table 1, we present specific numbers of cells in each population that responded to visual stimuli. A cell was categorized as responding if it increased its firing rate at the onset, during, or at the offset of light stimulation.

The common observation for all groups was a significant drop in the peak responses for each population (Figure 2). After suppression of photoreceptor activity by Sildenafil, all populations displayed a change in their response patterns. The Off-sustained (Figure 2A) group lost the excitatory component of the response and only reacted by suppressing the activity when the stimulus was on screen. The Off-Transient cells (Figure 2B) became On-Sustained-like, but with a delayed onset component. The On-Off group (Figure 2C) also lost the Off-response component. The patterns of the On-Sustained (Figure 2D), On-Transient (Figure 2E), and Other (Figure 2F) groups became almost identical to the Off-Transient and On-Off groups. Thus, most of the cells could no longer be assigned to their original group after losing photoreceptor activity. In all groups, we observed a dramatic drop in the activity evoked by a flash of light (Figure 2G), which was accompanied by a robust increase in response latency (Figure 2H). Moreover, we found that in the On and Other groups, the transient response lasted longer than under control conditions (Figure 2I), eventually becoming a persistent response.

**Figure 2.**
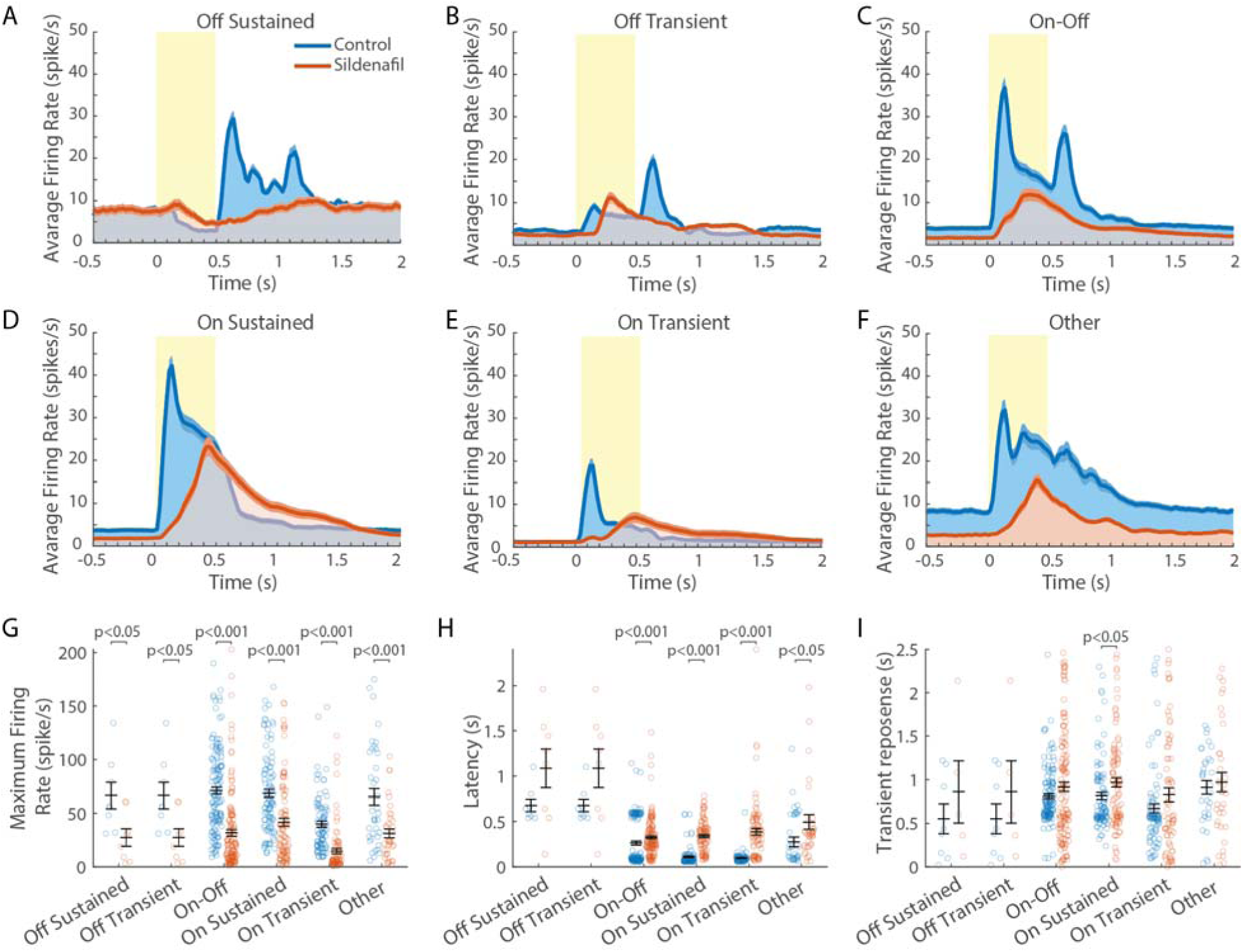
Silencing the phototransduction cascade changes the physiological profiles of the RGCs. Comparison of population histograms before (blue) and after Sildenafil application (red) for the following RGC types: A, Off-sustained cells; B, Off-transient cells; C, On-Off cells; D, On-sustained cells; E, On-transient cells; F, RGC cells that do not fit the classification of any of the above groups. G, Summary of the peak maximum firing rates before and after Sildenafil. H, Summary of latency change before and after Sildenafil. I, Duration of the prolonged response after stimulus offset, before and after Sildenafil.

Retinal ganglion cells have stimulus preferences similar to those in the Superior colliculus and primary visual cortex. To compare any changes in the stimulus selectivity, we recorded neuronal responses to different drifting-grating parameters before and after the addition of Sildenafil. Overall, in the control condition, we observed cells showing broad orientation selectivity, measured as the Half-Width at Half-Height (HWHH), broad spatial-frequency (SF) tuning, and broad temporal-frequency (TF) tuning. Broad SF and TF tuning means that cells can detect a large range of various parameters. This property changed dramatically after inhibition of phototransduction. Even though some cells kept their orientation selectivity despite a marked decrease in response, an entire population lost the ability to detect higher SFs (above 0.03 cycle/°) and TFs (above 1.5 cycle/s). Figure 3 shows examples of typical RGC physiology before (blue) and after Sildenafil (red), displayed as Orientation/Direction tuning curves (Figure 3A, B), SF tuning curves (Figure 3D, E), and TF tuning curves (Figure 3G, H). Comparison of the distribution of HWHH values did not show differences in population average (44.15 ± 20.14° *vs*. 45.23 ± 20.40°); however, there was a decrease in cell numbers after Sildenafil application (Figure 3C). That means the inhibition of photoreceptor activity didn’t change the orientation selectivity of the recorded population. The result was different in the case of the SF and TF values. As shown in the examples, the population almost entirely lost the ability to detect higher spatial frequencies, which resulted in an average population change from 0.082 ± 0.051 cycle/° to 0.038 ± 0.072 cycle/° (p < 0.005; Figure 3F). The difference was even more pronounced in the distribution of preferred TFs and population averages, where the change was from 3.5 cycle/s to 1 cycle/s (P < 0.001; Figure 3I). These results suggest that the recorded population kept the ability to detect orientation but lost visual acuity and fast temporal perception.

**Figure 3.**
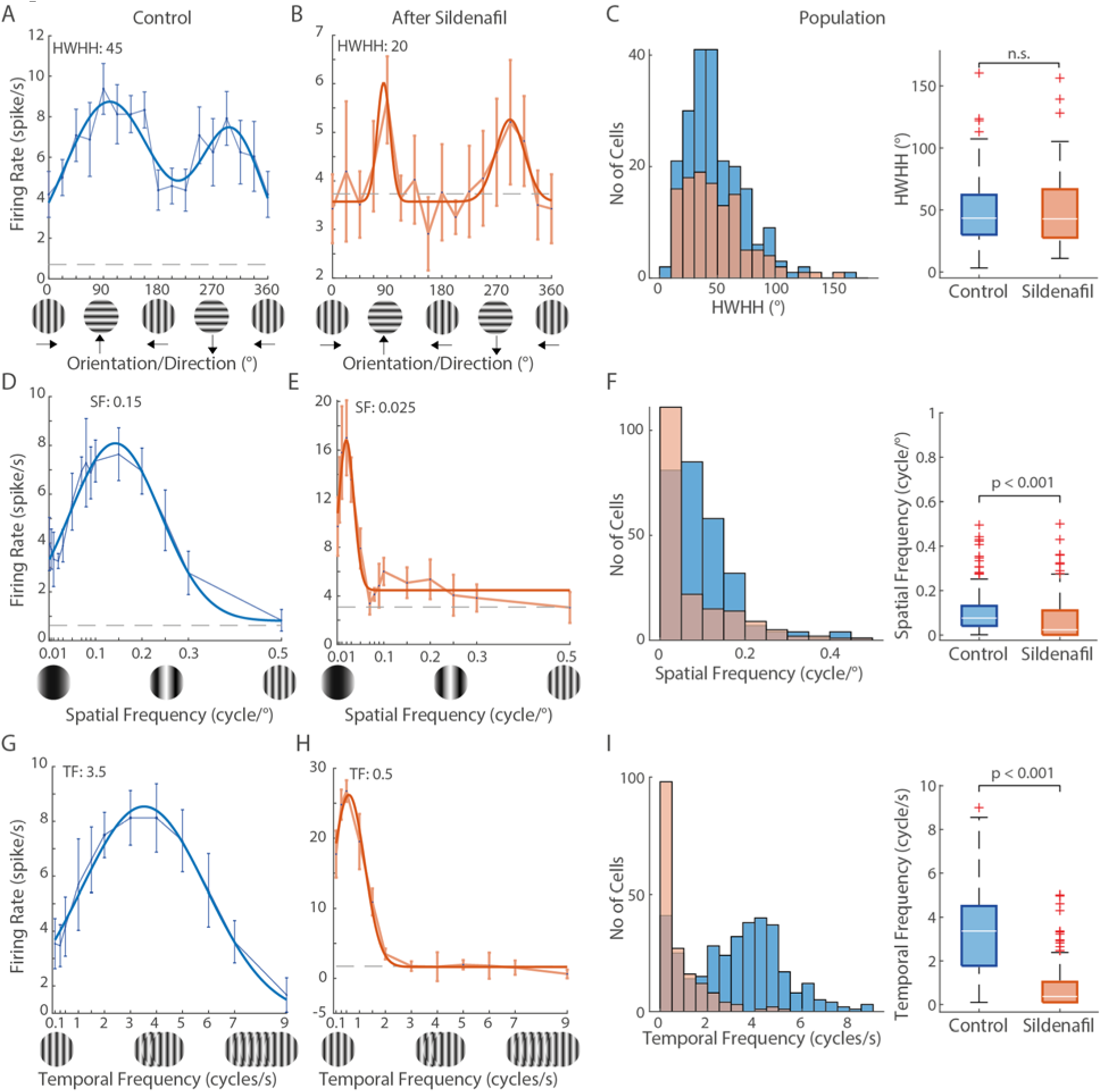
Spatiotemporal receptive field properties of RGCs before and after Sildenafil addition. A, Typical example of a control orientation-tuning curve. B, Typical example of an orientation-tuning curve after Sildenafil. C, Distribution of the HWHH values before and after Sildenafil, and corresponding summarizing whisker plots. D, Typical example of a control spatial-frequency tuning curve. E, Typical example of a spatial-frequency tuning curve after Sildenafil. F, Distribution of the preferred optimal SF values before and after Sildenafil, and corresponding summarizing whisker plots. G, Typical example of a control temporal-frequency tuning curve. H, Typical example of a temporal-frequency tuning curve after Sildenafil. I, Distribution of the preferred optimal TF values before (blue) and after Sildenafil (red), and corresponding summarizing whisker plots.

The change in physiology after PDE6 suppression was even more evident when we examined the behavior of individual cells. In Figure 4, we present four examples of RGCs whose electrophysiological profiles and parameter preferences are changed completely, revealing that some cells increased their firing rates after Sildenafil and developed stimulus selectivity. It was intriguing to see a subset of cells (*e.g.*, Cell 1 and 2; Figure 4A, B) that displayed the general trend of responses as in the entire population in the case of SF and TF, but showed different behavior in orientation tuning curves. A subset of cells showed an increased firing rate even if they lost the ability to detect higher spatial and temporal frequencies (Figure 4C, D). The most notable result was the transition of certain orientation-selective cells into direction-selective cells (all examples in Figure 4). The most curious examples are Cells 3 and 4; they either did not show any direction selectivity until after the addition of Sildenafil, or they increased their signal-to-noise ratio by changing from orientation to direction selectivity (Figure 4C, D). The raster plots in Figure 4, show the relation between Flash and Grating responses and changes of individual cells in response to different stimuli. The raster plots clearly indicate that most of the population (83% of cells for flash, 85% of cells for gratings) decreased their firing rate after Sildenafil in both types of stimuli; but a small part (10% of cells) showed the opposite behavior depending on stimulation (Figure 4E, F), displaying a change in the firing rate (after *vs*. before Sildenafil). We observed a similar pattern in background activity, with a somewhat higher percentage of cells with increased activity after Sildenafil (29% of cells had higher background activity, and 71% of cells had a lower background after Sildenafil, Figure 4G). To investigate in detail changes in the orientation selectivity, we considered only the subset of cells that preserved responses to drifting gratings after the addition of the PDE6 inhibitor. About half of those cells increased their Orientation-Selectivity Index (OSI; Figure 4H), and about the same number of cells showed increased Direction Index (DI; Figure 4I), suggesting that the same cells shifted toward more direction-selective cells with higher selectivity. On the other hand, quantification of the HWHH indicated only increased HWHH values (Figure 4J), which is related to decreased orientation selectivity. Finally, we extracted a population of cells that preserved responses to the tests of Spatial Frequency (Figure 4K) and Temporal Frequency (Figure 4L). As shown above, those cells mainly presented a decrease in preferred SFs and TFs.

**Figure 4.**
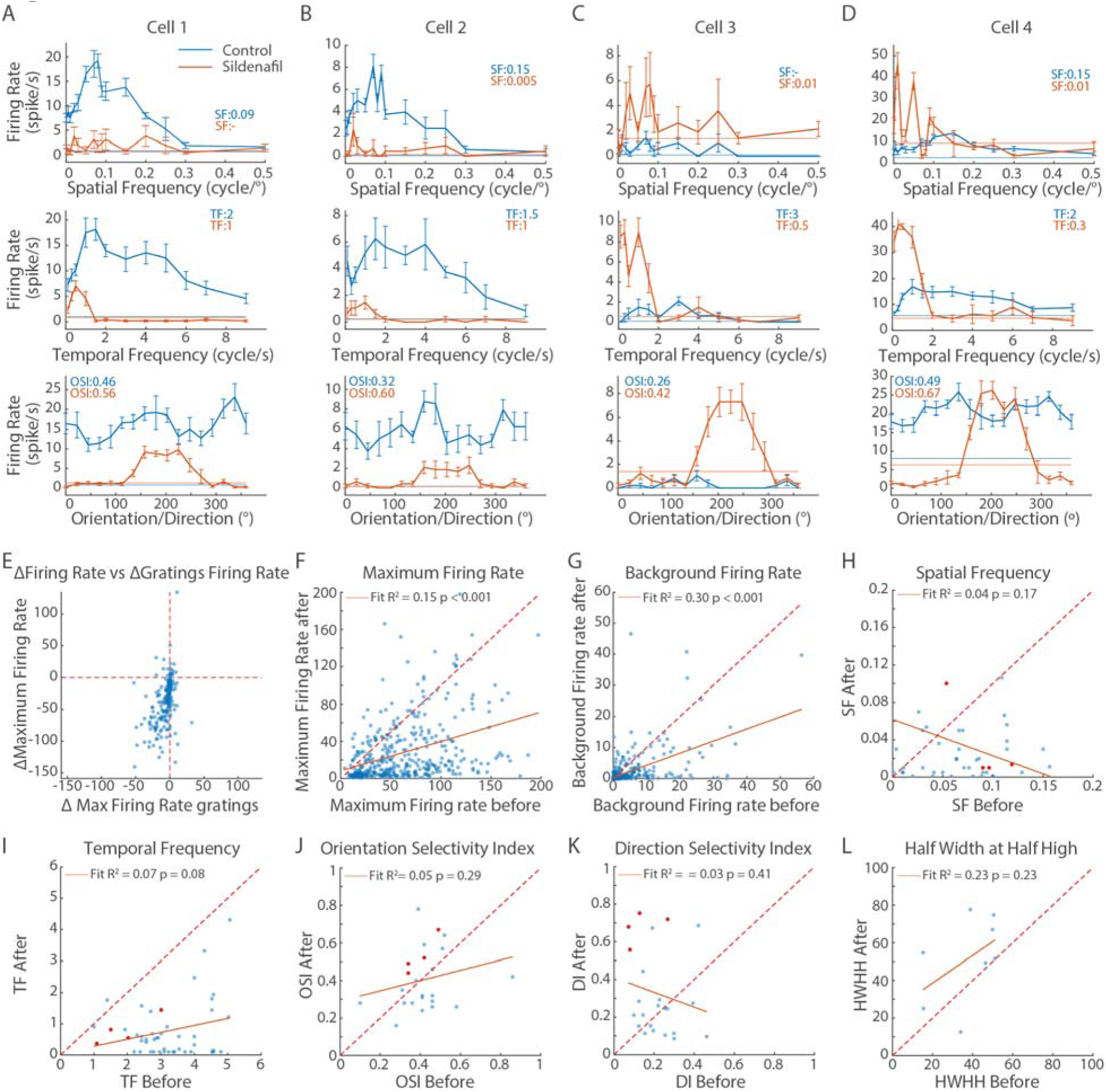
Reducing the amount of visually evoked input radically changes the properties of individual RGCs. A-D, Examples of cells that changed their stimulus selectivity after inhibition of photoreceptor activity by Sildenafil. E, Change in response (Firing Rate) to stimulation by flash (Y axis) and drifting gratings (X axis). F – G, Raster plots of activity, and properties of recorded neurons before (X axis) and after (Y axis) inhibition of photoreceptor activity by Sildenafil. Raster plots represent: F, maximum firing rate of response to flash; G, firing rate during spontaneous activity; H, corresponding orientation selectivity index (OSI) for the same cells; I, direction selectivity index (DI); J, orientation selectivity measured as half-width at half height (HWHH); K, spatial-frequency (SF) preference; L, temporal-frequency (TF) preference. Each dot represents a single cell. Red dots indicate cells shown in A-D. The solid lines indicate linear regression, characterized by correlation coefficients (R^2^ values), along with the correlation p-values as insets to raster plots. The dashed lines show a linear diagonal for better visualization of the data shift.

### Sildenafil effect is specific to cone photoreceptors

Our results created new questions related to this unexpected phenomenon. To answer the question of why we did not observe complete activity silencing, and to which neuronal pathway it is specific, we used two additional mouse models of retinal degeneration: *Gnat1^irdr^*(further called Gnat1) and *Gnat2^cpfl3^* (further called Gnat2), to turn off rod or cones, respectively. The effect of sildenafil in both models was robust and clear. Basic analyses of VEPs before and after Sildenafil implementation in Gnat1 and Gnat2 revealed that the process is more selective to cone photoreceptors; however, it has a partial effect on rod photoreceptors as well (Figure 5A-D). The effect is presented in Figure 5A, where a flat line represents a lack of visually evoked responses in Gnat1 mice when Sildenafil was applied. This is also confirmed in population data analysis of the VEP amplitudes and VEP response latencies. The difference between the two models was made more prominent when looking at the flash response histograms, which were one of the main features of Sildenafil’s influence on the retina (Figure 5E-H). In Gnat1 mice, where only cones were active, the addition of Sildenafil completely removed visually evoked responses (Figure 5E), unlike in the case of Gnat2, where only rods were active (Figure 5F). What was notable the Gnat2 responses after Sildenafil looked very similar to healthy retinas after inhibitor treatment. This clearly indicates the Sildenafil selectivity towards the cone PDE6. Because there were no visually evoked responses in Gnat1 mice after Sildenafil, we only compared drifting grating responses of Gnat2 and regular C57B6 mice after Sildenafil application to see if we observed a similar effect (Figure 5I-P; Table 2). Generally, the RGC properties of the Gnat2 mutant mouse did not differ in parameter distribution from the healthy animals. However, there were specific differences in visual information processing after Sildenafil application (Figure I-L). The distribution of orientation tuning widths shifted towards narrower responses compared with both untreated Gnat2 mice and treated C57B6 mice (Figure 5M). We did not observe such a difference between treated and non-treated C57B6 mice. Another interesting finding was the lack of a shift towards very low SF after the Sildenafil treatment in the Gnat2 mouse model (Figure 5N). The preference slightly shifted towards broader gratings; however, unlike in healthy mice, it did not saturate in the lowest tested SF. There was a strong similarity in the TF processing after Sildenafil treatment (Figure 5O). Both in C57B6 and Gnat2 mice retinas became slow frequencies band-pass filter, meaning we observed detection only of very slow stimuli (Figure 5P).

**Figure 5.**
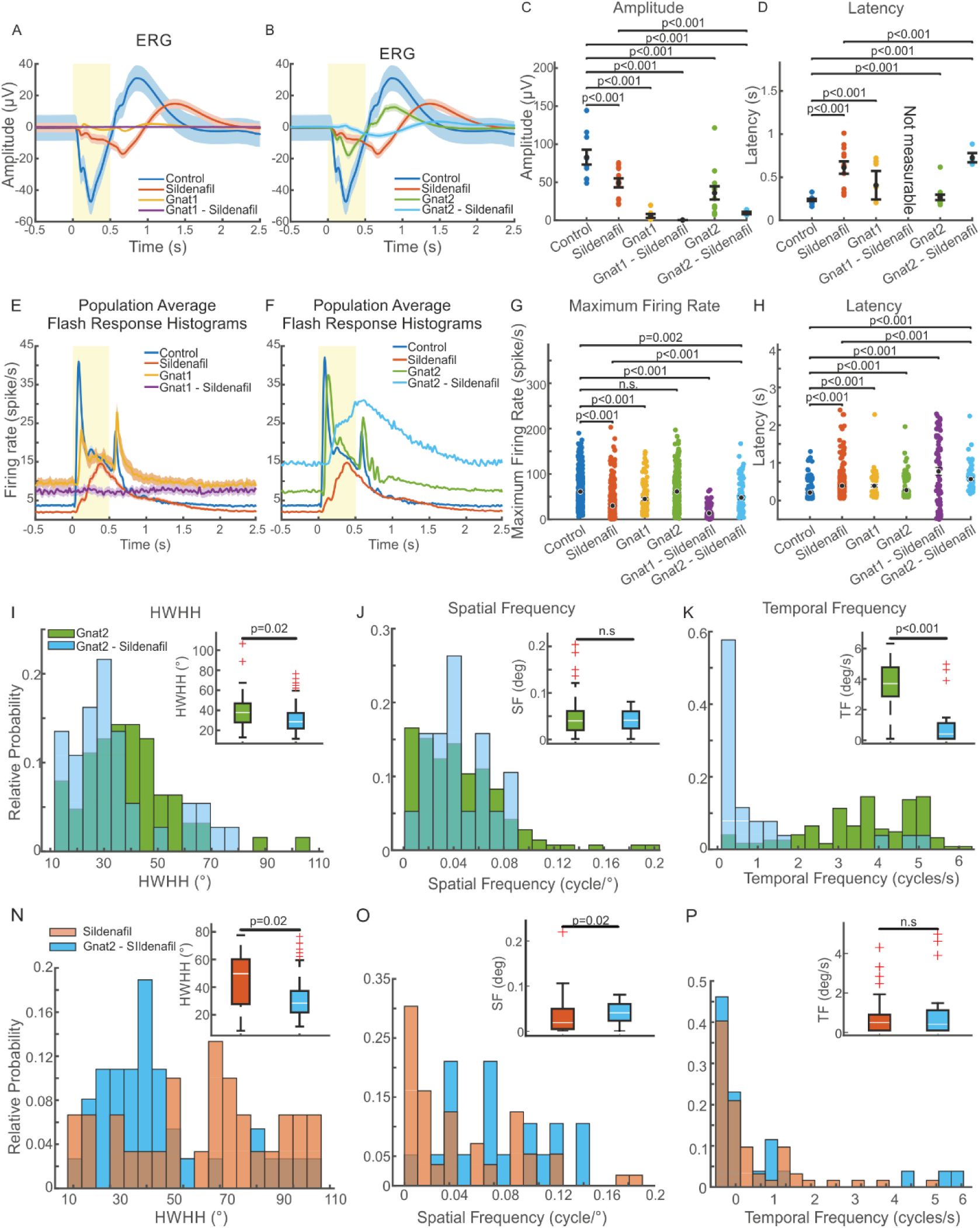
Specificity of Sildenafil to cone photoreceptor indicated by electrophysiology. A, the ERGs for Gnat1 and C57B6 mice before and after treatment. B, similar responses as in A, but for Gnat2 and C57B6 mice. C, population summary of ERG amplitudes before and after treatment for all 3 models used. D, population summary of the response latencies for all 3 mouse models. E, Gnat1, and C57B6 mice population responses to flash stimulus. F, Gnat2, and C57B6 mice population response histograms to flash stimulus. G, population summary of all 3 mouse models and their maximum response intensity before and after Sildenafil treatment. H, similar summary as in G, but indicating response latency. I, tuning the width of population distributions before and after treatment for Gnat2 mice. J, population distribution of optimal spatial frequencies found in Gnat2 mice before and after treatment. K, population distributions of the optimal temporal frequencies for Gnat2 mice as in J. L, whisker plots summarizing all three parameters shown in I–K. M, distribution comparison of tuning width after Sildenafil treatment for C57B6 (red) and Gnat2 (blue) mice. N, similar comparison as in M, but for the optimal spatial frequencies. O, similar comparison as in M and N, but for optimal temporal frequencies. P, the general summary and comparison of tuning width, SFs, and TFs for C47B6 and Gnat2 mice after PDE6 inhibitor application.

**Table 2.** Drifting gratings parameter comparison between mouse models.

|  | Half Width Half<br>Height | Spatial<br>Frequency | Temporal<br>Frequency |
| --- | --- | --- | --- |
| Control | 44.15 ± 1.87 | 0.082 ± 0.01 | 3.53 ± 0.08 |
| Sildenafil | 45.23 ± 3.72 | 0.038 ± 0.01 | 1 ± 0.22 |
| Gnat1 | 47.97 ± 4.01 | 0.06 ± 0.01 | 3.19 ± 0.17 |
| Gnat2 | 39.26 ± 2.16 | 0.04 ± 0.003 | 3.58 ± 0.13 |
| Gnat2-Sildenafil | 33.2 ± 2.83 | 0.04 ± 0.01 | 0.92 ± 0.27 |

## Discussion

Suppressing the phototransduction cascade has an enormous impact on retinal information processing in animals (Barabás et al., 2003; Zhang et al., 2005; Wang et al., 2011; Niell, 2013; Kinoshita et al., 2015; Cote, 2021) and humans (Glossmann et al., 1999; Laties and Zrenner, 2002a; Papageorgiou et al., 2018; Karaarslan, 2020; Ausó et al., 2021). Our data indicate that decreasing photoreceptor activity completely changes the spatial and temporal preference of Retinal Ganglion Cells (RGCs), and the On or Off properties of distinct cell types. Many studies have already shown how Sildenafil affects the On and Off responses of particular RGC types (Martins et al., 2015; Østergaard et al., 2020). Our results confirm a drop in the evoked activity of RGCs, which is the most prominent effect. However, we have also observed that most groups of cells, except for one, changed to On-like cell types with long response latencies. The Off-Sustained cells preserved partial suppression of the activity when the light was on; however, like the Off-Transient and On-Off populations, they totally lost the Off-response component. When comparing response histograms, five of the six groups that we characterized displayed almost the same activity and properties profiles after the addition of Sildenafil, except for slight differences in response latency. The Off-response loss suggests the strongest impact on the Off-processing pathway when photoreceptors are suppressed. At the moment, we do not know whether this phenomenon occurs via a disinhibitory mechanism due to insufficient input to inhibitory cells (for example, horizontal or AII amacrine cells) or *via* decreased input to rod bipolar cells (Marc et al., 2007). The aspect that should be considered here is the relative affinity of Sildenafil for the various isoforms of the PDE enzyme, and their distribution in the various cell types. In this regard, it has been reported that Sildenafil has a stronger affinity to the cone PDE6 variant (Nivison-Smith et al., 2014; Cote, 2021).

Another interesting question emerges, namely, whether our results reflect electrophysiological changes that occur during photoreceptor degeneration. There is not much evidence about detailed visual processing in models of retinal degeneration because most of the literature focuses on flash responses (Fransen et al., 2015; Carleton and Oesch, 2024; Dyszkant et al., 2025). Considering the basic parameters of the responses (Yu et al., 2017), such as firing-rate change or latency, our results align with changes previously found in degenerated retinas *ex vivo* (Allen et al., 2010; Borowska et al., 2011; Brown et al., 2011; Dyszkant et al., 2025). However, it is much harder to resolve changes in the orientation/direction selectivity and spatial or temporal preference (Vaney et al., 2012; Ding et al., 2016; Wei, 2018; Dhande et al., 2019; Kerschensteiner, 2022; Kim et al., 2022), which occur after blocking the PDE6. As Sildenafil has a stronger affinity for cone-photoreceptor PDE6, it would be logical to conclude that our findings should resemble features of the visual responses displayed by mouse models of rod (GNAT1) or cone (GNAT2) dysfunctions (Wang et al., 2011). Our recorded spatial and temporal single-cell preferred parameters, however, are indicative of different phenomena. Thus, this hypothesis requires more data, because the visually evoked responses in the GNAT1 and GNAT2 models would be much different and would require very thorough analysis. There is a possibility of remnant responses, as suggested by the findings with double GNAT-knock-out mice, where VEPs were recorded intracranially (Flood et al., 2022); those responses could come from melanopsin-expressing ganglion cells. However, no information exists on the tuned responses mediated by the melanopsin pathway.

Our results demonstrate that Sildenafil’s action on visual processing has a higher affinity toward the cone’s PDE6. This was confirmed by comparing *Gnat1* (rod-deficient, cone-only) and *Gnat2* (cone-deficient, rod-only) mouse models. In *Gnat1* mice, where only cones remain functional, the addition of Sildenafil completely abolished visually evoked responses, consistent with a direct block of cone phototransduction. In contrast, in *Gnat2* mice, where rods provide the sole photoreceptor input, responses persisted and were similar to those of wild-type retinas after Sildenafil treatment. These findings strongly support prior biochemical studies reporting a higher affinity of Sildenafil for cone PDE6 isoforms (Zhang et al., 2005; Cote, 2021) and extend them by providing functional electrophysiological evidence at the level of retinal ganglion cell spatiotemporal tuning. The cone selectivity also explains why high-frequency spatial and temporal responses – characteristic of cone-driven pathways – were abolished after PDE6 inhibition. Moreover, the selective suppression of cone input provides a mechanistic basis for the transient visual side effects reported in humans after PDE5/6 inhibitor use, including impaired flicker sensitivity, altered color vision, and photophobia (Laties and Zrenner, 2002b; Ausó et al., 2021). Beyond pharmacology, these findings establish Sildenafil application as a pharmacological tool to transiently silence cone pathways in otherwise intact retinas, creating a model system to study inner retinal adaptation to cone loss and to mimic aspects of early cone dysfunction in degenerative diseases.

To explain the shift in temporal- and spatial-frequency preference of single ganglion cells, we need more experiments dedicated to specific transmission or connectivity to broaden current knowledge and extend this research, as well as in various degenerative models, which would also help to understand how the network physiology changes during photoreceptor degeneration.

## Acknowledgments

We are grateful to all the Ophthalmic Biology Group members for their insights about this project and for providing a supportive environment.

The International Centre for Translational Eye Research (FENG.02.01-IP.05-T005/23) project is carried out within the International Research Agendas programme of the Foundation for Polish Science co co-financed by the European Union under the European Funds for Smart Economy 2021-2027 (FENG) National Science Center, Poland (2019/34/E/NZ5/00434) for ATF National Science Center, Poland (2022/47/B/NZ5/03023) for ATF

## Conflict of interest

The authors declare no competing financial interests.

## Notes

### Competing Interest Statement

The authors have declared no competing interest.

